# Chromosomal-level reference genome of the incense tree *Aquilaria sinensis*

**DOI:** 10.1101/2020.03.03.972679

**Authors:** Wenyan Nong, Sean T.S. Law, Annette Y.P. Wong, Tobias Baril, Thomas Swale, Lee Man Chu, Alexander Hayward, David T.W. Lau, Jerome H.L. Hui

## Abstract

Trees in the genus *Aquilaria* (Thymelaeaceae) are known as lign aloes, and are native to the forests of southeast Asia. Lign aloes produce agarwood as an antimicrobial defence. Agarwood has a long history of cultural and medicinal use, and is of considerable commercial value. However, due to habitat destruction and over collection, lign aloes are threatened in the wild. We present a chromosomal-level assembly for *Aquilaria sinensis*, a lign aloe endemic to China known as the incense tree, based on Illumina short-read, 10X Genomics linked-read, and Hi-C sequencing data. Our 783.8Mbp *A. sinensis* genome assembly is of high physical contiguity, with a scaffold N50 of 87.6Mbp, and high completeness, with a 95.8% BUSCO score for eudicotyledon genes. We include 17 transcriptomes from various plant tissues, providing a total of 35,965 gene models. We reveal the first complete set of genes involved in sesquiterpenoid production, plant defence, and agarwood production for the genus *Aquilaria*, including genes involved in the biosynthesis of sesquiterpenoids via the mevalonic acid (MVA), 1-deoxy-D-xylulose-5-phosphate (DXP), and methylerythritol phosphate (MEP) pathways. We perform a detailed repeat content analysis, revealing that transposable elements account for ∼61% of the genome, with major contributions from *gypsy*-like and *copia*-like LTR retroelements. We also provide a comparative analysis of repeat content across sequenced species in the order Malvales. Our study reveals the first chromosomal-level genome assembly for a tree in the genus *Aquilaria* and provides an unprecedented opportunity to address a variety of applied, genomic and evolutionary questions in the Thymelaeaceae more widely.

## Introduction

The genus *Aquilaria* (family Thymelaeaceae) contains fifteen tree species, commonly known as ‘lign aloes’, native to the forests of southeast Asia. A special feature of lign aloes is their production of agarwood, which is also known as aloeswood or gharuwood. Agarwood is produced as an antimicrobial defence mechanism, after infection of the tree with a fungal pathogen, and involves the saturation of infected heartwood with a dark aromatic resin.

The main active compounds present in agarwood are terpenoids, specifically sesquiterpenes and derivatives of flindersiachromone (Chen et al. 2012), and the composition of oil extracted from agarwood is exceedingly complex, including over 150 chemical compounds (Naef 2011). As a consequence of the unique fragrant properties of agarwood, it has long been traded as a highly prized cultural, religious, and medical commodity. For example, the use of agarwood as a fragrant product is recorded in Sanskrit Vedas dating to 1,400 B.C., while the Greek physician Pedanius Dioscorides recorded medical uses for agarwood in his *De Materia Medica* from 65 A.D., and agarwood is also highly revered as an icense in Islamic, Buddhist and Hindu ceremonies (Lopez-Sampson and Page 2018).

Today, the demand for agarwood remains great, not least in the form of oud oil which is distilled from agarwood for perfumery, and high grade agarwood products can reach prices as high as US$100,000/kg, with global trade estimated at US$6-US$8 billion (Akter et al. 2013). A driver of the expense of agarwood products is the depletion of wild lign aloes, which has led to their inclusion on Appendix II of the Convention on International Trade in Endangered Species of Wild Fauna and Flora (CITES), in an attempt to control and monitor international trade and help limit impact, while species in the genus have been categorized as “vulnerable” and “critically endangered” by the IUCN Red List of Threatened Species (Newton and Soehartono 2001; Harvey-Brown 2018).

The incense tree, *Aquilaria sinensis* (Lour.) Spreng, is a lign aloe endemic to the south eastern part of China (530,906 km^2^, Harvey-Brown 2018), including Hainan Province, Fujian, Guanxi, Guangdong Provinces, and Hong Kong. The tree is relatively slow growing, taking 50 to 100 years to reach its maximum height of ∼15 meters (CITES 2015). Owing to the overexploitation of natural populations, *A. sinensis* is currently included on the “List of Wild Plant Under State Protection (Category II)” in mainland China (The State Council, People’s Republic of China 1999), while it is protected under different ordinances in Hong Kong (Agriculture, Fisheries and Conservation Department 2018). In response to the conservation challenges facing *A. sinensis*, plantations have been initiated, including >10,000 hectares of land in south eastern China from 2006 to 2013 (Yin et al. 2016), and annual planting of 10,000 seedlings in the country park and other suitable habitats in Hong Kong (Agriculture, Fisheries and Conservation Department 2018).

In addition to its ecological and scientific importance, *A. sinensis* holds particular cultural significance in Hong Kong, as it is commonly believed that the species provided the region’s name, which translates from Chinese as ‘Incense Harbour’ or ‘Fragrant Harbour’. Meanwhile, the trading of agarwood or *Chen Xiang/Cham Heong* (translated from Mandarin/Cantonese) was an important industry since the Sung Dynasty (610-970) (Iu 1983; Agriculture, Fisheries and Conservation Department 2003). Although the cultivation of *A. sinensis* for the agarwood industry in Hong Kong ceased during in the last century, remaining populations continue to persist in the countryside of Hong Kong, including lowland and broad-leaved forests (Yip and Lai 2004).

Despite the great scientific and cultural importance of *A. sinensis*, a high-quality genome is lacking, hindering further understanding of the species, and scientifically driven conservation measures. More widely, genomic resources for the genus *Aquilaria* are poor, with only a chloroplast genome of *A. sinesis* and a draft genome of *A. agallocha* available currently (Chen et al. 2014; Wang et al. 2016). To address this issue, here we provide a high quality chromosomal-level genome assembly for *A. sinesis* together with a large number of accompanying transcriptomes from diverse plant tissues.

## Materials and Methods

### Sample collection and Genome sequencing

Tissue samples used for genome sequencing (mature leaf) and transcriptome sequencing (young and mature leaves, young shoot, flower buds, flower, fruit, seed, aril, intact and wounded stem) were collected from incense tree individuals on the campus of The Chinese University of Hong Kong during the flowering and fruiting period in June 2019. During sample collection, healthy leaves without visible symptoms of fungal infection (e.g. leaf spot, rust or wilt) were collected for DNA extraction. Both surfaces of the leaves were cleaned and rinsed with double-distilled water prior to DNA extraction to minimise the potential contamination. Genomic DNA (gDNA) was extracted from *A. sinensis* using a DNeasy® Plant Mini Kit (Qiagen) following the manufacturer’s protocol. Extracted gDNA were subjected to quality control using Nanodrop spectrophotometer (Thermo Scientific) and gel electrophoresis. High molecular DNA was extracted with CTAB method and had a mean molecular weight of 85 kbp. Qualifying samples were sent to Novogene, and Dovetail Genomics for library preparation and sequencing. The resulting library was sequenced on Illumina HiSeq□X platform to produce 2□×□150 paired-end sequences. The length-weighted mean molecule length is 23590.15 bases, and the raw data can be found at NCBI’s Small Read Archive (SRR10737433). Details of the sequencing data can be found in Supplementary information S1.

### Chicago library preparation and sequencing

A Chicago library was prepared as described previously (Putnam et al, 2016). Briefly, ∼500ng of HMW gDNA (mean fragment length = 85 kbp) was reconstituted into chromatin *in vitro* and fixed with formaldehyde. Fixed chromatin was digested with DpnII, the 5’ overhangs filled in with biotinylated nucleotides, and free blunt ends were ligated. After ligation, crosslinks were reversed and the DNA purified from protein. Purified DNA was treated to remove biotin that was not internal to ligated fragments. The DNA was then sheared to ∼350 bp mean fragment size and sequencing libraries were generated using NEBNext Ultra enzymes and Illumina-compatible adapters. Biotin-containing fragments were isolated using streptavidin beads before PCR enrichment of each library. The libraries were sequenced on an Illumina HiSeq X to produce 211 million 2×150 bp paired end reads, which provided 146.61 x physical coverage of the genome (1-100 kb pairs).

### Dovetail HiC library preparation and sequencing

A Dovetail HiC library was prepared in a similar manner to that described previously (Erez Lieberman-Aiden et al., 2009). Briefly, for each library, chromatin was fixed in place with formaldehyde in the nucleus and then extracted fixed chromatin was digested with the restriction enzyme DpnII, the 5’ overhangs were filled in with biotinylated nucleotides, and free blunt ends were ligated. After ligation, crosslinks were reversed and the DNA purified from protein. Purified DNA was treated to remove biotin that was not internal to ligated fragments. The DNA was sheared to ∼350 bp mean fragment size and sequencing libraries were generated using NEBNext Ultra enzymes and Illumina-compatible adapters. Biotin-containing fragments were isolated using streptavidin beads before PCR enrichment of each library. The libraries were sequenced on an Illumina HiSeq X sequencer to produce 193 million 2×150 bp paired end reads, which provided 66,743.81 x physical coverage of the genome (10-10,000 kb pairs).

### Transcriptome sequencing

Transcriptomes of multiple developmental stages were sequenced at Novogene (Figure 2, Supplementary information S2). Total RNA from different tissues were isolated using a combination method of cetyltrimethylammonium bromide (CTAB) pre-treatment (Jordon-Thaden et. al. 2015) and mirVana™ miRNA Isolation Kit (Ambion) following the manufacturer’s instructions. The extracted total RNA was subjected to quality control using a Nanodrop spectrophotometer (Thermo Scientific), gel electrophoresis, and an Agilent 2100 Bioanalyzer (Agilent RNA 6000 Nano Kit). Qualifying samples underwent library construction and sequencing at Novogene; polyA-selected RNA-Sequencing libraries were prepared using TruSeq RNA Sample Prep Kit v2. Insert sizes and library concentrations of final libraries were determined using an Agilent 2100 bioanalyzer instrument (Agilent DNA 1000 Reagents) and real-time quantitative PCR (TaqMan Probe) respectively. Details of the sequencing data can be found in Supplementary Information S1.

**Figure 1.**
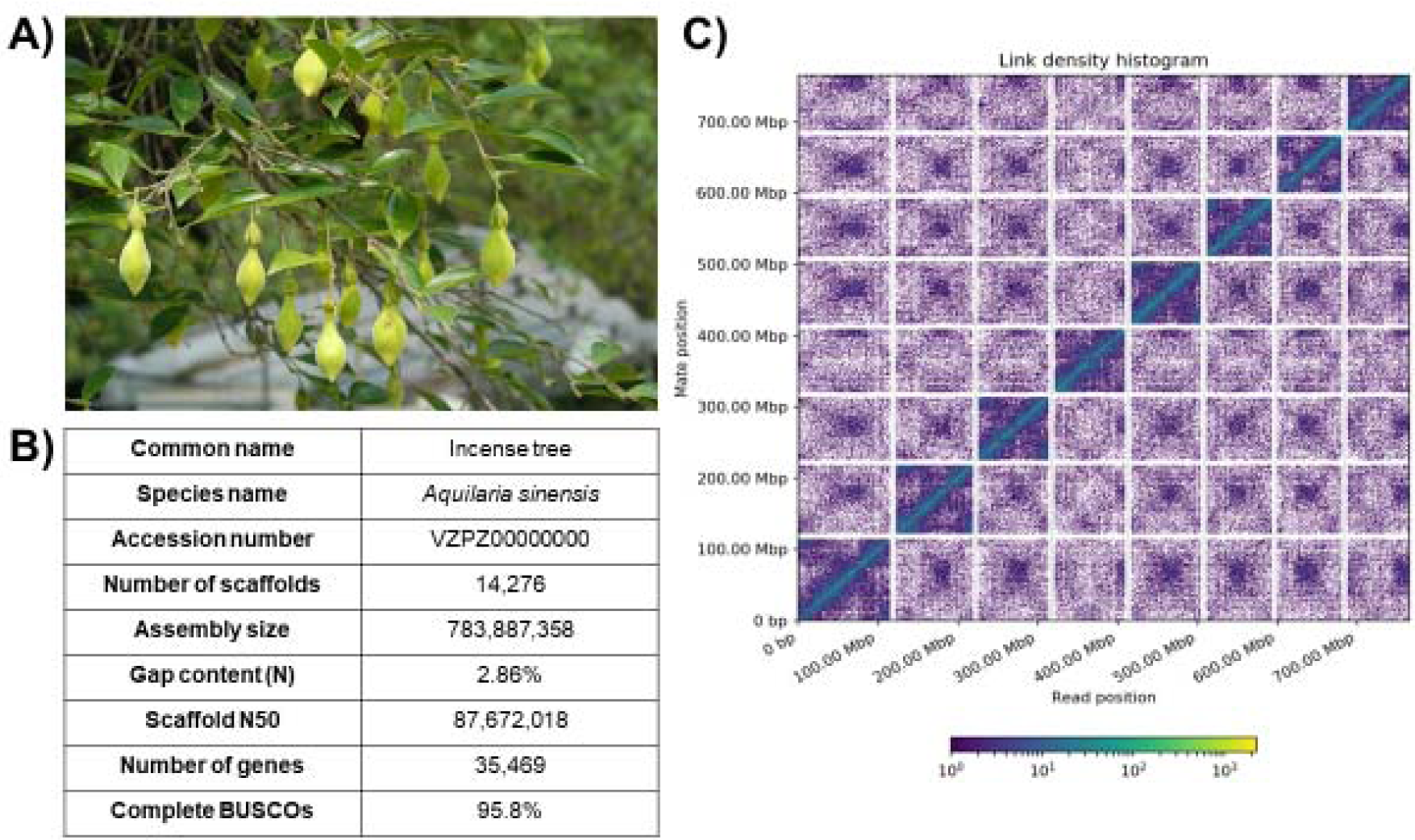
**A) Photograph of an incense tree *Aquilaria sinensis* (Lour.) Spreng taken in Hong Kong; B) *Aquilaria sinensis* Hi-C information.** The x- and y- axes provide the mapping positions for the first and second reads in each read pair respectively, grouped into bins. The colour of each square indicates the number of read pairs within that bin. White vertical and black horizontal lines have been added to indicate the borders between scaffolds. Scaffolds less than 1 Mb are excluded.

**Figure 2.**
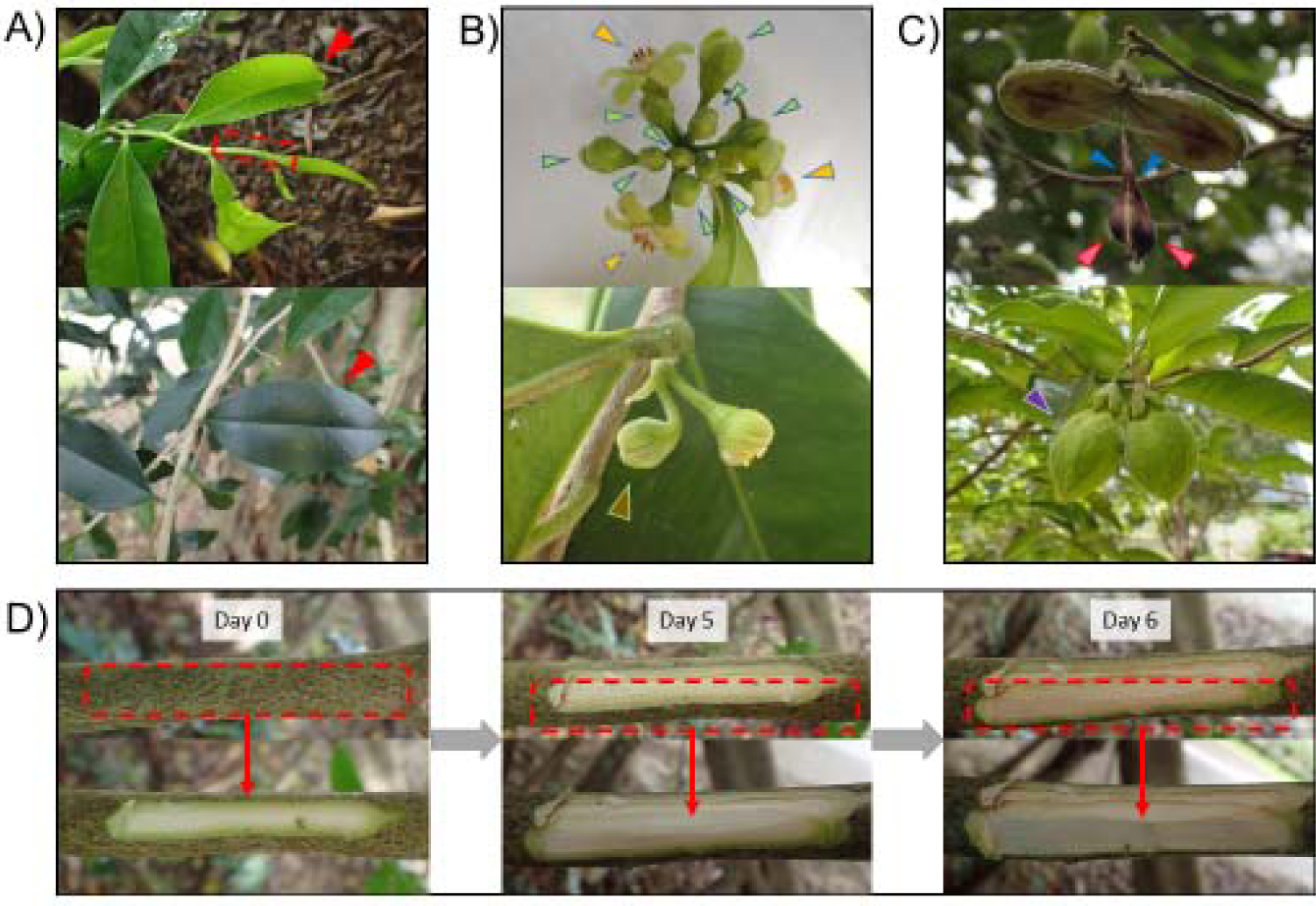
Photographs of tissues from different developmental stages used for transcriptome sequencing. **A) Young and mature leaves and a young shoot used for transcriptome sequencing**. Tissues used for RNA extraction are indicated in red**; B) Floral tissues used for transcriptome sequencing**. Flower, flower bud and fertilized flower are indicated in yellow, green and brown respectively. **C) Fruit tissues used for transcriptome sequencing**. Fruit arils, seeds and flesh (pericarp and mesocarp) are indicated in blue, pink and purple. **D) Intact and wounded stem tissues used for transcriptome sequencing**. Photos showing before and after the extraction of stem tissues (approximately 5 cm x 1 cm, indicated in red) from intact (day 0) to wounded (day 5 and 6) conditions.

### Genome assembly

Linked-Read data were assembled using the Supernova™ assembler (v2.1.1, Zheng et al 2016; Marks et al 2019), using the default recommended settings (https://support.10xgenomics.com/de-novo-assembly/software/overview/latest/performance) to produce the a pseudohaplotype assembly outputs (--style=pseudohap). The Supernova output pseudohaplotype assembly, shotgun reads, Chicago library reads, and Dovetail HiC library reads were used as input data for HiRise, a software pipeline designed specifically for scaffolding genome assemblies using proximity ligation data (Putnam et al 2016). An iterative analysis was conducted as follows. First, Shotgun and Chicago library sequences were aligned to the draft input assembly using a modified SNAP read mapper (http://snap.cs.berkeley.edu). The separations of Chicago read pairs mapped within draft scaffolds were analyzed by HiRise to produce a likelihood model for genomic distance between read pairs, and the model was used to identify and break putative misjoins, to score prospective joins, and make joins above a threshold. After aligning and scaffolding Chicago data, Dovetail HiC library sequences were aligned and scaffolded following the same method. After scaffolding, shotgun sequences were used to close gaps between contigs.

### Gene model prediction

Raw sequencing reads from 17 transcriptomes were pre-processed with quality trimmed by trimmomatic (v0.33 with parameters “ILLUMINACLIP:TruSeq3-PE.fa:2:30:10 SLIDINGWINDOW:4:5 LEADING:5 TRAILING:5 MINLEN:25”, Bolger et al. 2014). For the nuclear genomes, the genome sequences were cleaned and masked by Funannotate (v1.6.0, https://github.com/nextgenusfs/funannotate) (Palmer and Stajich 2017), the softmasked assembly were used to run “funannotate train” with parameters “--stranded RF -- max_intronlen 350000” to align RNA-seq data, ran Trinity, and then ran PASA (Haas et al 2008a). The PASA gene models were used to train Augustus in “funannotate predict” step following recommended options for eukaryotic genomes (https://funannotate.readthedocs.io/en/latest/tutorials.html#non-fungal-genomes-higher-eukaryotes). Briefly, the gene models were predicted by funannotate predict using the following parameters: “--repeats2evm --protein_evidence uniprot_sprot.fasta -- genemark_mode ET --busco_seed_species embryophyta --optimize_augustus --busco_db embryophyta --organism other --max_intronlen 350000”. The gene models originated from several prediction sources, including: ‘GeneMark (Lomsadze et al 2005)’: 76877, ‘HiQ’: 12640, ‘pasa’: 26175, ‘Augustus (Stanke et al 2006)’: 18455, ‘GlimmerHMM (Majoros et al, 2003)’: 189664, ‘snap (Korf 2004)’: 120842, ‘total’: 444653’. Gene models were passed to Evidence Modeler (Haas et al 2008b) (EVM Weights: {‘GeneMark’: 1, ‘HiQ’: 2, ‘pasa’: 6, ‘proteins’: 1, ‘Augustus’: 1, ‘GlimmerHMM’: 1, ‘snap’: 1, ‘transcripts’: 1}) to generate the final annotation files, and PASA (Haas et al 2008b) was used to update the EVM consensus predictions, add UTR annotations and models for alternatively spliced isoforms.

### Repetitive elements annotation

Repetitive elements were identified using an in-house pipeline. Firstly, elements were identified using RepeatMasker ver. 4.0.8 (Smit et al 2013) with the *eukarya* RepBase repeat library (Jurka et al 2005). Low-complexity repeats were not masked and the sensitive search parameter was specified. Following this, a *de novo* repeat library was constructed using RepeatModeler ver. 1.0.11 (Smit and Hubley 2015), including RECON ver. 1.08 (Bao and Eddy 2002) and RepeatScout ver. 1.0.5 (Price et al 2005). Novel repeats identified by RepeatModeler were analysed using a ‘BLAST, Extract, Extend’ process to improve the characterisation of elements along their entire length (Platt et al 2016). Consensus sequences were generated for each family, along with classification information. The resulting *de novo* repeat library was utilised to identify repetitive elements using RepeatMasker. Longer repeats were constructed using RepeatCraft (Wong et al 2018) using LTR_FINDER ver. 1.0.5 (Xu et al 2007) to defragment repeat segments. At loci where RepeatMasker annotations overlapped (i.e where the same sequence was annotated as different repeat families), only the longest repeat was kept. This conservative approach helps avoid TE content estimates being inflated by counting the same bases multiple times and ensures a one-to-one matching of sequence with repeat family identity. A revised summary table was constructed with the final repeat counts. All plots were generated using Rstudio ver. 1.2.1335 (Racine and Rstudio 2015) with R ver. 3.5.1 (Team 2013) and ggplot2 ver. 3.2.1 (Wickham 2016).

### Gene family annotation and gene tree building

Potential gene family sequences involved in the mevalonic pathway, methylerythritol phosphate pathway, and jasmonic acid biosynthesis pathway were first identified by homology searching in the KEGG database, and were retrieved from the genome using tBLASTn. Identity of each putatively identified gene was then tested by comparison to sequences in the NCBI nr database using BLASTx. For phylogenetic analyses of gene families, DNA sequences were translated into amino acid sequences and aligned to other members of the gene family; gapped sites were removed from alignments using MEGA and phylogenetic trees were constructed using MEGA (Kumar et al 2018).

## Results and Discussion

### High quality Aquilaria genome

Genomic DNA was extracted from single individuals of *Aquilaria sinensis* (Figure 1A), and sequenced using the Illumina short-read and 10X Genomics linked-read sequencing platforms (Supplementary information S1). Hi-C libraries were also constructed and sequenced on the Illumina platform. The genome sequences were first assembled with short-reads followed by scaffolding with Hi-C data. The genome assembly is 783.8 Mbp with a scaffold N50 of 87.6 Mbp (Table 1). This high physical contiguity is matched by high completeness, with a 95.8% % complete BUSCO score for eudicotyledon genes (version odb10) (Table 1). A total of 35,965 gene models using 17 transcriptomes from tissues collected from different developmental stages generated in this study were incorporated (Figure 2, Supplementary information S1), with mean exon length being 304 bp, mean intron length being 518 bp, and mean deduced protein length being 338 aa. The majority of the sequences assembled (∼91%) were contained on 8 pseudomolecules (Figure 1B, supplementary information S2), representing the first near chromosomal-level genome generated for species in the genus *Aquilaria*. This high-quality *Aquilaria* genome provides an unprecedented opportunity to address the issue of a variety of applied, genomic and evolutionary questions in the Thymelaeaceae more widely.

**Table 1.**
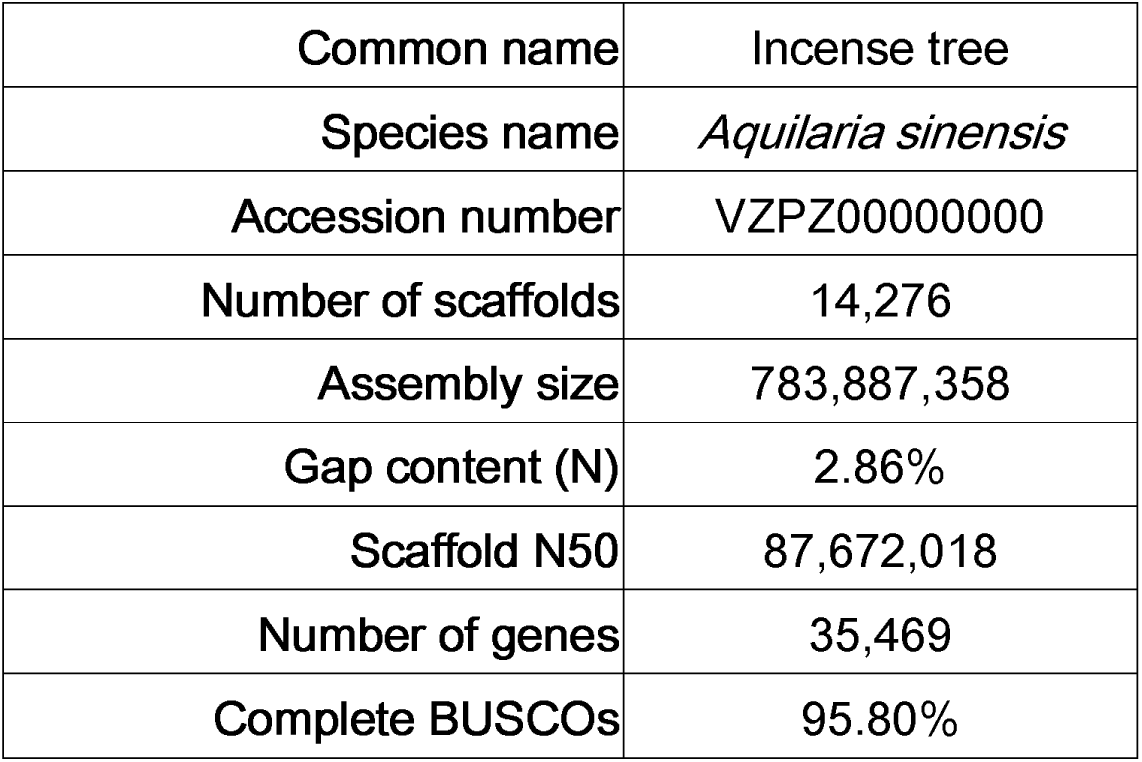
Table summarising genome assembly statistics.

### Transposable elements

Transposable elements (TEs) make up a proportion of most eukaryote genomes, but the contribution of TEs to genome size varies greatly across lineages. In plants, TEs may dominate the host genome, for example comprising 85% of the maize genome (Schnable et al. 2009) and potentially as much as 90% of the crown imperial (*Fritillaria imperialis*) genome (Ambrožová et al. 2010). (Ambrožová et al. 2010). At the other end of the spectrum, TEs may account for a very small proportion of the genome, such as in the carnivorous bladderwort, where TEs represent just 3% of total genomic DNA (Ibarra-Laclette et al. 2013). However, in most plant genomes TE content lies somewhere between these extremes.

We estimate that the repeat content of the incense tree accounts for more than half of its total genome size (61.22%)(Figure 3A, Supplementary information S3), demonstrating that TE dynamics have played an important role in shaping the genome evolution of this species. Of the repeat types present in the incense tree genome, very few were annotated as small RNA, satellite, simple or low complexity repeats (∼0.9% of the total genome), with the majority of the genome (∼61%) consisting of TEs (i.e. SINEs, LINEs, LTR retrotransposons, DNA transposons)(Figure 3A, Supplementary information S3). By far the main contribution of TEs to the incense tree genome is from LTR elements, which account for over one third of the total assembly (36.47%, Figure 3A, Supplementary information S3). DNA transposons represent 6.4% of the incense tree genome, and there is just a small contribution from LINEs (2.1% of the genome), and very few SINEs present in the genome (0.01%)(Figure 3A, Supplementary information S3).

**Figure 3.**
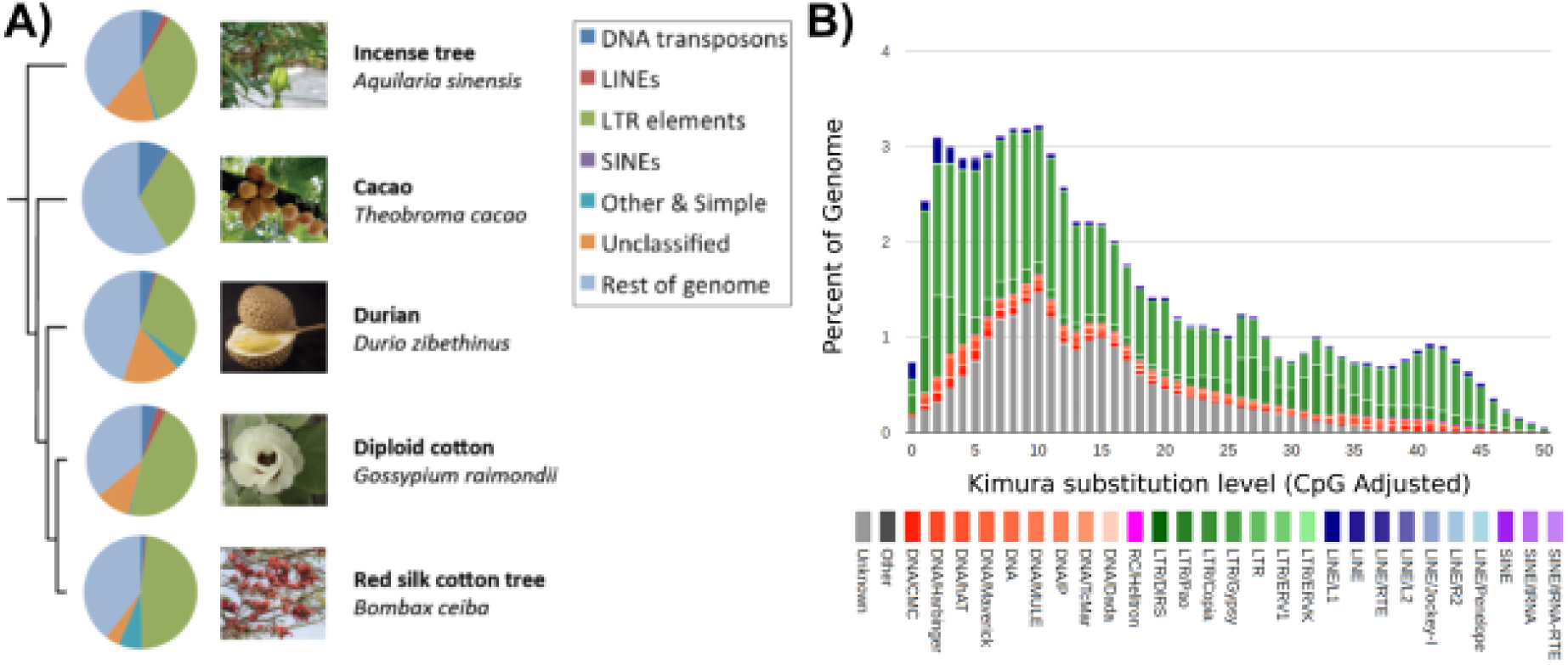
**A) Comparison of repeat content among Malvales genomes.** The phylogeny indicates evolutionary relationships among species in the Malvales with available genome assemblies. Pie charts illustrate the proportions of different repeat types in each genome, as indicated by the colours presented in the key. The incense tree photograph is provided under a GNU Free Documentation License to Wikipedia user Chong Fat (https://en.wikipedia.org/wiki/File:HK_Aquilaria_sinensis_Fruits.JPG), the durian photograph is provided under a GNU Free Documentation License to Wikipedia user 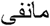 (https://en.wikipedia.org/wiki/File:Durian_in_black.jpg), the cacao photograph is provided under a GNU Free Documentation License to Wikipedia user Adolfoovalles (https://en.wikipedia.org/wiki/File:Matadecacao.jpg), the red silk cotton tree photograph is provided under a GNU Free Documentation License to Wikipedia user Lonely Explorer (https://en.wikipedia.org/wiki/File:Bombax_ceiba_-_Cotton_tree_-_Shimul_Flower_(2).jpg), and the diploid cotton photograph was taken from the USDA National Cotton Germplasm Collection (NCGC) (https://www.cottongen.org/organism/Gossypium/raimondii). For the cacao genome, ‘other’, ‘simple’ and ‘unclassified’ repeat types were not reported, reflecting the differences in relation to the other genomes. **B) Transposable element (TE) accumulation history in the incense tree genome**, based on a Kimura distance-based copy divergence analysis of TEs, with Kimura substitution level (CpG adjusted) illustrated on the x-axis, and the percentage of the genome represented by each repeat type on the y-axis. Repeat type is indicated by the colour chart below the x-axis.

In Figure 3A, we summarise recently reported TE analyses for sequenced members of the order, specifically: diploid cotton (Wang et al 2012), red silk cotton tree (Gao et al 2018), durian (The et al 2017), and cacao (Matina 1-6) (Motamayor et al 2013). This provides a comparative overview of TE content in species closely related to the incense tree. Differences in the sequencing strategies employed to generate available Malvales genomes, accompanying variation in assembly quality, and differences in the approaches applied to generate TE annotations all complicate comparative analysis of TEs among members of the Malvales. However, it is clear that the incense tree shows the same general trends as other species in the order (Figure 3A). In summary, TEs make up a large proportion of total genomic content across the Malvales, with LTR retrotransposons representing by far the greatest contribution to repeat content (32-48%, Supplementary information S4). As with other members of the Malvales, *gypsy*-like and *copia*-like LTR elements were the most abundant superfamilies among the LTR elements identified in the incense tree genome (The et al 2017; Motamayor et al 2013)(Figure 3A, Supplementary information S4). DNA transposons make up the next largest contribution to Malvales genomes, but they represent a considerably lower proportion of the genome (1-9%) (Figure 3A, Supplementary information S4). Among DNA transposons, MULE elements were the most abundant superfamily, followed by En/Spm elements from the CMC superfamily (Supplementary information S4). LINEs make up a very small proportion of total TE content (0.17-2.7%), with SINEs almost completely absent from Malvales genomes (0.09% in *G. raimondii*, 0.01% in *A. sinensis*) (Figure 3A). Overall, the observed patterns demonstrate that TEs represent a major force for shaping genome size among members of the Malvales, with LTR elements being by far the most important repeat class.

A Kimura distance-based copy divergence analysis for the incense tree revealed that the most frequent TE sequence divergence relative to TE consensus sequence was 1-10%, mainly due to increased distances among LTR elements (and unclassified elements), suggesting a relatively recent burst of sustained activity, followed by a gradual decline (Figure 3B).

### Genes involved in plant defence and agarwood production

Formation of agarwood is usually associated with fungal infection or physical wounding, where resin composed of mixtures of sesquiterpenes and 2-(2-phenylethyl)chromones (PECs) are secreted by the tree as a defence mechanism (Naef 2011; Chen et al 2012). Over time, the accumulation of volatile compounds and sesquiterpenoids lead to the formation of fragrant agarwood (Fazila and Halim 2012; Hashim et al 2014).

In the *A. sinensis* genome, genes involved in the biosynthesis of sesquiterpenoids from isoprenoid precursors using the mevalonic acid (MVA) pathway in the cytosol and the 1-deoxy-D-xylulose-5-phosphate (DXP) or methylerythritol phosphate (MEP) pathway in the plastid were all found to be present (Figure 4, supplementary information S5). Further, these two pathways biosynthesise C5 homoallylic isoprenoid precursors including isopentenyl pyrophosphate (IPP) and dimethylallyl pyrophosphate (DMAPP), and the genes encoding enzymes for production of IPP and DMAPP could all be identified (Figure 4, supplementary information S5). These C5 isoprene units are later converted to C10 geranyl pyrophosphate (GPP) and C15 farnesyl pyrophosphate (FPP) in the presence of the key-limiting enzymes GPP synthase and FPP synthase (Figure 4, supplementary information S5). In the final stage of sesquiterpenes production, the necessary sesquiterpene synthase enzymes (SesTPS) can also be identified in 74 loci (Figure 4, supplementary information S5 and S6). Further, the jasmonic acid (JA) signalling pathway has been reported to be involved in plant defence and it plays a significant role in the wound-induced signalling mechanism in regulating SesTPS expression (Xu et al. 2017; Tan et al. 2019), and genes involved in its biosynthesis pathway could all be identified (Figure 4, supplementary information S5).

**Figure 4.**
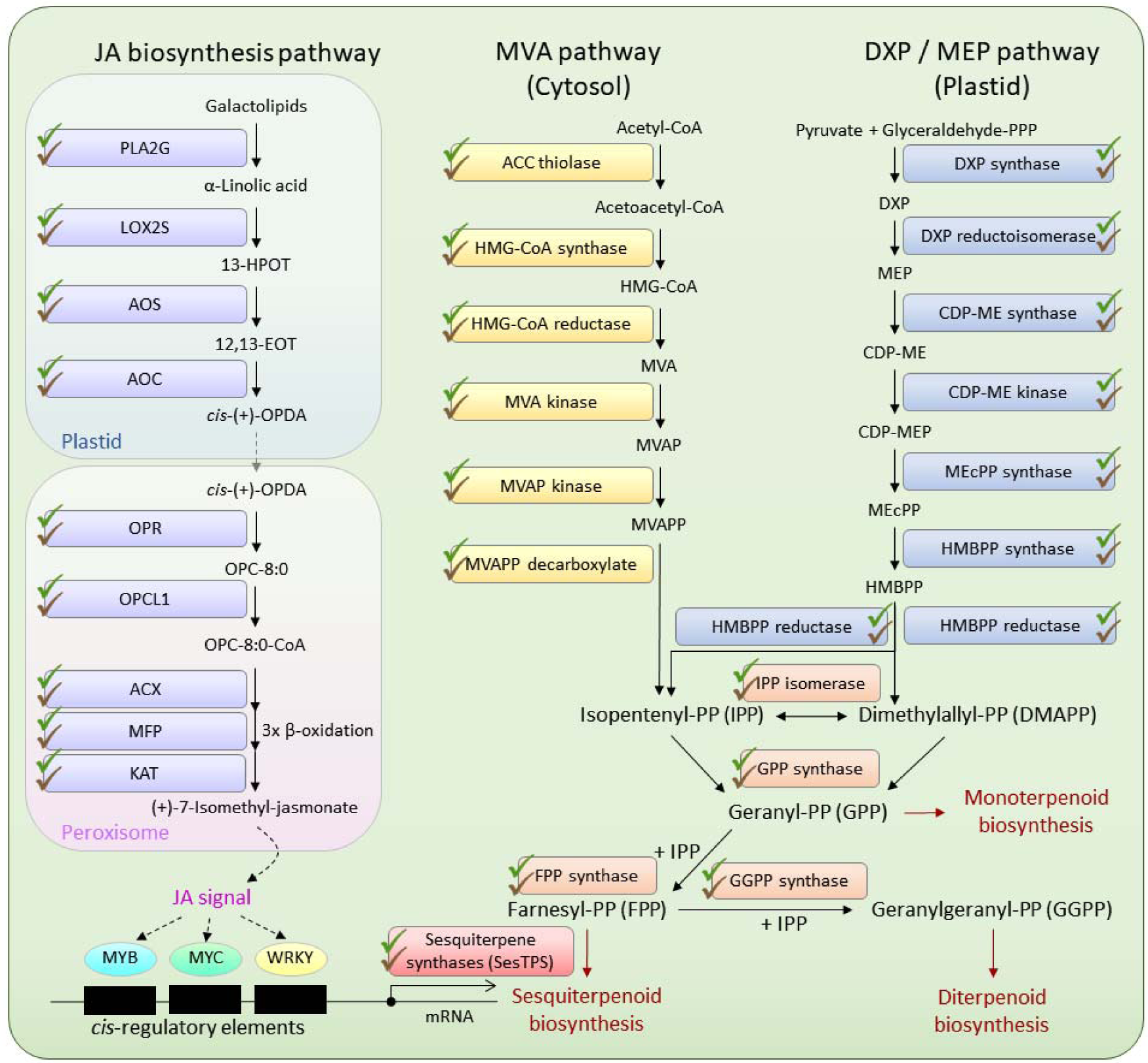
Genes involved in plant defence and agarwood production in *Aquilaria sinensis*, including the Mevalonic acid (MVA) pathway, 1-deoxy-D-xylulose-5-phosphate (DXP) or methylerythritol phosphate (MEP) pathway and jasmonic acid (JA) biosynthesis pathway. Green ticks represent genes that can be identified in the genome, and brown ticks depict genes that can also be identified in the wounded stem transcriptome data. Abbreviations: (MVA pathway) ACC, Acetyl-CoA; HMG-CoA, hydroxymethylglutaryl-CoA; MVAP, phosphomevalonate; MVAPP, diphosphomevalonate; (DXP/MEP pathway) CDP-ME, 4-diphosphocytidyl-2-C-methyl-D-erythritol; MEcPP, 2-C-Methyl-D-erythritol-2,4-cyclopyrophosphate; HMBPP, (E)-4-Hydroxy-3-methyl-but-2-enyl pyrophosphate; (JA biosynthesis pathway) PLA2G, secretory phospholipase A2; LOX2S, lipoxygenase; 13-HOPT, 13(S)-hydroperoxy-octadecatrienoic acid; AOS, allene oxide synthase; 12,13-EOT, 12,13(S)-epoxy-octadecatrienoic acid; AOC, allene oxide cyclase; *cis*-(+)-OPDA, cis-(+)-12-oxo-phytodienoic acid; OPR, 12-oxophytodienoic acid reductase; OPC-8:0, 3-oxo-2-(2′(Z)-pentenyl)-cyclopentane-1-octanoic acid; OPCL1, OPC-8:0 CoA ligase 1; ACX, acyl-CoA oxidase; MFP, multifunctional protein; KAT, 3-ketoacyl-CoA thiolase.

Interestingly, we have also found that genes from the all of the aforementioned pathways, including the MVA pathway, DXP or MEP pathway, and JA biosynthesis pathway, are all expressed in at least one of the wounded stem transcriptomic data sets (stem 02 and 03) (Figure 4). These data represent the first complete set of genes documented to be involved in sesquiterpenoid production, plant defence, and agarwood production in a single species of *Aquilaria*.

## Conclusion

This study presents the first high-quality genome assembled for a plant in the genus *Aquilaria*. The *A. sinensis* genome holds important scientific, commercial and conservation relevance, and our work provides details of the first complete set of genes involved in sesquiterpenoid and agarwood production in *Aquilaria*. More broadly, this high quality *A. sinesis* genome provides a useful reference point for further understanding of Thymelaeaceae biology and evolution.

## Acknowledgements

This study was supported by The Chinese University of Hong Kong Direct Grant (4053248), and the Agriculture, Fisheries and Conservation Department, The Government of the Hong Kong Special Administrative Region (AFCD/SQ/60/18) (JHLH). AH is supported by a Biotechnology and Biological Sciences Research Council (BBSRC) David Phillips Fellowship (BB/N020146/1). TB is supported by a studentship from the Biotechnology and Biological Sciences Research Council-funded South West Biosciences Doctoral Training Partnership (BB/M009122/1). STSL is supported by a postgraduate studentship from The Chinese University of Hong Kong.

## Data Accessibility

The raw genome and RNA sequencing data have been deposited in the SRA under Bioproject numbers SRR10737433 and PRJNA534170. The final chromosome assembly was submitted to NCBI Assembly under accession number VZPZ00000000 in NCBI.

## Author Contributions

JHLH conceived and supervised the study. WN, SLTS, AYPW, TS, TB, AH, JHLH performed the genome assembly, gene model prediction, gene annotation, and analyses. WN, STSL, AYPW, TS, TB, AH, LMC, DTWL, JHLH wrote the manuscript.

## Table, Figure and Supplementary Information

**Supplementary information S1**. Summary of genomic and transcriptomic data generated in this study.

**Supplementary information S2**. Statistics of the eight assembled pseudomolecules.

**Supplementary information S3**. Summary of repeat types present in the genome of the incense tree.

**Supplementary information S4**. Summary of repeat types present in the genomes of the incense tree and other members of the Malvales for which genopne assemblies exist.

**Supplementary information S5**. Gene sequences and locations of genes identified in Figure 4.

**Supplementary information S6**. A) Neighbour-joining tree of sesquiterpene synthases; B) Maximum-likelihood tree of sesquiterpene synthases.

## Notes

https://www.ncbi.nlm.nih.gov/bioproject/PRJNA534170

