## Supplementary information S3 for "Chromosomal-level reference genome of the incense tree *Aquilaria sinensis*"

Supplementary Table S1. Repeat content summary for the incense tree (*Aquilaria sinensis*), produced using a combination of library-based and de novo repeat analyses

| **Summary Genome Information** | |
| --- | --- |
| Sequences in Query Fasta | 15,358 |
| Total Genome Length (bp) | 787,906,105 (765,481,005 excl N/X-runs) |
| GC level | 37.98% |
| Bases masked | 482,392,546 (61.22%) |

| **Class/Family** | **Number of Elements** | **Length Occupied (bp)** | **Percentage of Sequence (%)** |
| --- | --- | --- | --- |
| **SINEs** | 467 | 71,806 | 0.01 |
| **LINEs** | 25,299 | 16,662,662 | 2.11 |
| **LTR elements** | 246,922 | 287,346,678 | 36.47 |
| **DNA elements** | 107,352 | 50,383,707 | 6.39 |
| **Unclassified** | 224,446 | 120,834,671 | 15.34 |
| **Total interspersed repeats** | 604,486 | 475,299,524 | 60.32 |
| **Other (Simple, Small RNA, Satellites, Low Complexity)** | 16,902 | 7,093,022 | 0.90 |
